# Cell enlargement causes mitotic errors and aneuploidy in cells that evade senescence after CDK4/6 inhibition

**DOI:** 10.1101/2025.07.01.662622

**Authors:** Aanchal U Pareri, Marialucrezia Losito, Floris Foijer, Adrian T Saurin

## Abstract

CDK4/6 inhibitors (CDK4/6i) arrest the cell cycle in G1 leading to cellular overgrowth and p53-dependent senescence. They are used to treat metastatic HR+/HER2-breast cancer, but resistance is common, and this has been associated with TP53 loss and senescence evasion. We show here that enlarged CDK4/6i-treated cells that evade senescence mis-segregate chromosomes due to defective chromosomal alignment and a weakened mitotic checkpoint, leading to aneuploidy and DNA damage. The chromosome alignment errors are associated with impaired Sgo1 localisation to centromeres and defective sister chromatin cohesion during mitosis. Importantly, all these mitotic defects can be rescued by constraining cell size during the CDK4/6i-treatment, and specifically restoring cohesion rescues the chromosome segregation errors. Together, this demonstrates mechanistically how cell enlargement drives genetic and karyotypic change in cells that re-enter the cell cycle following CDK4/6 inhibition. This could help fuel the rapid emergence of chemotherapy-resistant clones, especially in p53-null cells that evade senescence to drive drug-resistance in patients.

---

CDK4/6 inhibitors (CDK4/6i) are game-changing drugs that have revolutionised breast cancer treatment^1,2^. They arrest the cell cycle in G1 by preventing retinoblastoma (Rb) phosphorylation to inhibit E2F transcription factors and suppress S-phase entry^3–5^. During this G1 arrest, oncogenic signals in cancer cells activate mTOR to induce constitutive cell growth in the absence of proliferation, leading to cellular enlargement and permanent cell cycle exit into senescence^6–9^. This processes has been termed geroconversion^10^, and it is associated with a range of cellular stresses that promote senescence entry. Cellular enlargement in the absence of DNA replication induces a starvation-like remodelling of the proteome^11–15^, leading to cytoplasmic dilution^15^, osmotic stress^14^ and p38/p53/p21-dependent senescence from G1^6,14,16^. These enlarged cells have a weakened DNA damage response^16^ and if they fail to enter senescence from G1 they re-enter the cell cycle when the drug is removed and experience replication stress and further p21 induction during G2^14,17^. Similar effects were observed following inhibition of CDK2^6^ or CDK7^18^, which also arrest the cell cycle in G1, implying that geroconversion into senescence is a common outcome following a prolonged G1 arrest.

A major unresolved issue concerns the fate of cells that manage to evade senescence following CDK4/6i treatment. This occurs frequently in p53 deficient cells because these cells cannot induce p21 in response to the stresses that result from cell enlargement^6,14,16,17^. A high proportion of enlarged p53-deficient cells then re-enter the cell cycle and reach mitosis, were they experience chromosome segregation errors, leading to gross nuclear abnormalities and further DNA damage^6,14,17^. These damaged cells are also able to continue proliferating, due to p53-loss and defective DDR signalling, which we hypothesise drives genetic and chromosomal instability (CIN) and the rapid emergence of drug-resistant clones. In support, various clinical studies have shown that TP53 is rarely mutated in patients that respond well to CDK4/6i-treatment, but it is frequently mutated in cases of intrinsic or acquired resistance^19–22^. Furthermore, a very recent clinical study by Kudo et al links CDK4/6i-resistance to a lack of p21 induction in TP53-null cells, which prevents geroconversion into senescence^19^. It is therefore crucial to understand why enlarged CDK4/6i-treated cells that evade senescence missegregate chromosomes because the resulting CIN is likely to fuel the acquisition of drug-resistant karyotypes and genotypes.

We show here that enlarged cells that fail to enter senescence following CDK4/6i-treatment have defective centromeric cohesion and a weakened mitotic checkpoint, which combine to cause chromosome segregation errors. These defects result from the increased cell size which weakens the mitotic checkpoint and redistributes Sgo1 away from centromeres, leading to premature loss of cohesion. Together, this reveals how increases in cell size can dysregulate key mitotic processes to drive aneuploidy and genomic damage, with important implications for the evolution of CDK4/6i-resisitance in breast cancer.

## Results

### Cellular enlargement induces chromosome segregation errors and aneuploidy

CDK4/6i-treatment increases cell size and causes severe chromosome segregation errors when enlarged cells reach mitosis after drug removal^6,14,17^. The increased cell size causes the mitotic errors because these are abolished by restricting overgrowth using direct or indirect mTOR inhibition during the cell cycle arrest^6,14,17^. To study mechanistically why cell enlargement causes mitotic defects we compared cells that had been arrested with palbociclib (hereafter CDK4/6i) for different lengths of time in the presence or absence of PF-05212384 (hereafter mTORi) which prevents cell overgrowth^6,14^ (Figures S1A). We analysed non-transformed RPE1 cells and the HR+/HER2-breast cancer line, MCF7, which both arrest efficiently and overgrow extensively in G1 following CDK4/6i-treatment^6,14,16^. All drugs were then washed out and the cells analysed as they progressed through the first mitosis (Fig.1A).

**Figure 1.**
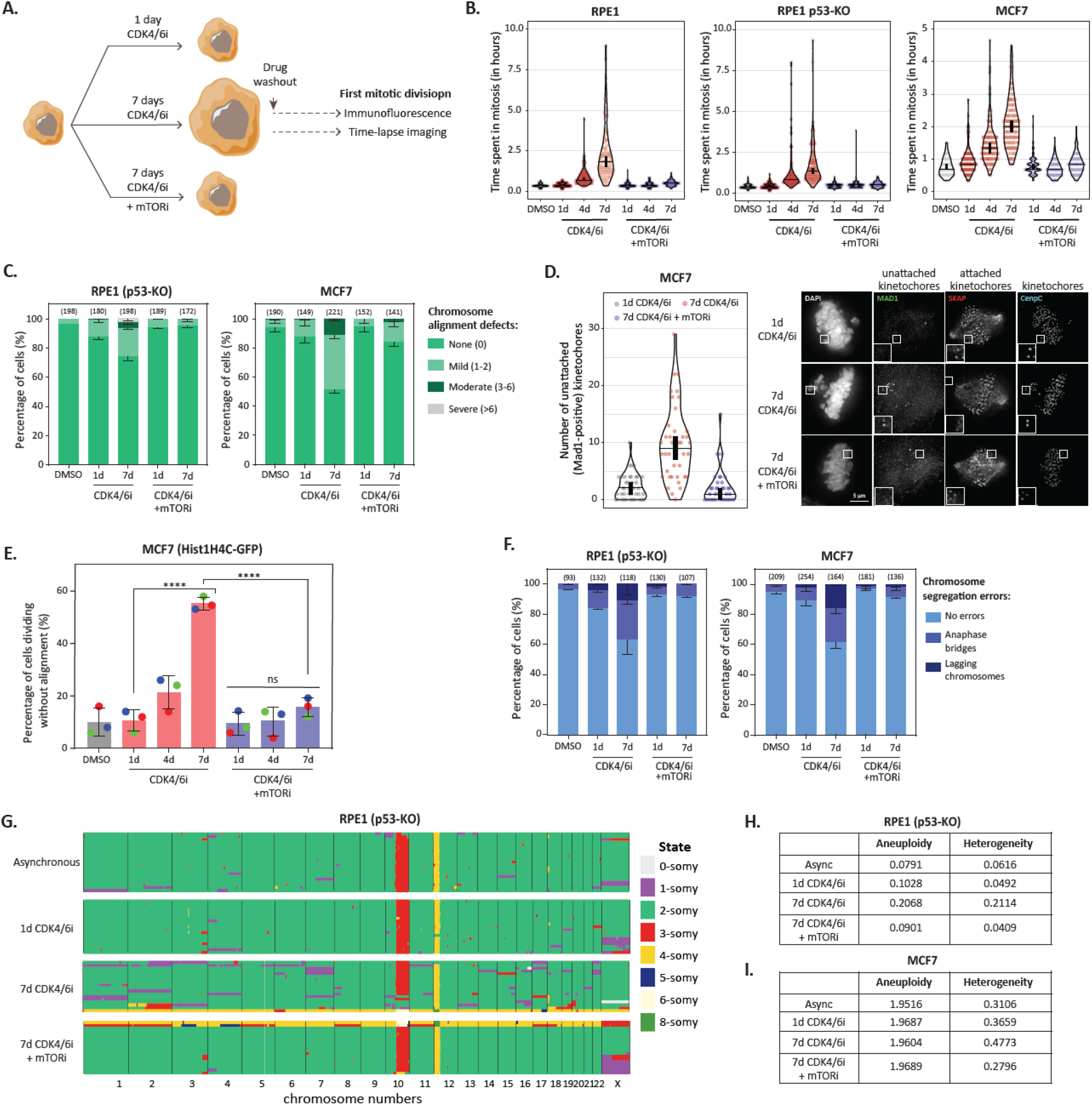
Enlarged CDK4/6i-treated cells experience a range of mitotic defects after drug washout. **A)** Schematic showing the treatment protocols to arrest cell in G1 with/without increasing cell size. Time lapse imaging or immunofluorescence was then performed during the first mitotic event after drug washout. **(B-F)** Effects of G1 arrest duration and cell size on: B) Mitotic duration (100-150 cells from 3 experiments); C) Percentage of chromosome misalignments (± SEM; 3 experiments, with total number of cells quantified per condition indicated above each bar); D) Number of unattached kinetochores per cell, based on MAD1 positivity (30 cells from 3 experiments with MG132 included to prevent cells exiting mitosis). Right panel shows example images of unattached kinetochores; E) Percentage cells dividing before chromosome alignment is complete. Statistical significance was determined by Chi-square test (****p<0.0001). Data shows mean ± SEM from three repeats; F) Percentage of chromosome segregation errors (± SEM; 3 experiments, with total number of cells quantified per condition indicated above each bar). **(G)** Heat maps showing copy number variations in RPE1 p53-KO treated as indicated before releasing for 72h prior to single cell sequencing. 24 single cell clones analysed per condition.

Live cell imaging demonstrated that enlarged RPE1 and MCF7 cells that recover from a prolonged G1 arrest experience significant delays in the first mitosis (Figure 1B). A lower proportion of enlarged RPE1 cells reach mitosis after drug washout, which is known to be due to earlier entry into senescence from G1 or G2^6,14,17^. This can be bypassed by knockout of p53, which allows the RPE1 cells to synchronously enter mitosis similarly to MCF7 cells (Figure S1B). We next sought to characterise the mitotic defects in these enlarged cells that evade senescence following CDK4/6i treatment. Immunofluorescence imaging demonstrated that the mitotic delays are associated with chromosome alignment errors and an inability to attach chromosomes to microtubules via the kinetochore (Figure 1C-D). However, these cells still exited mitosis prematurely with misalignments, leading to lagging chromosomes and anaphase bridges (Figure 1E-F). All these mitotic defects are size-dependent because they are rescued by inhibiting mTOR during the G1 arrest to prevent cell enlargement (Figs.1B-F & S1). This shows that a prolonged G1 arrest causes later defects in chromosome segregation during mitosis. These defects do not result for the prolonged G1 arrest *per se*, but rather they result from continue cell growth that occurs during that arrest.

This has important clinical implications because cells that manage to evade senescence following CDK4/6i-treatment, for example p53-deficient cells, could use their enlarged size to become chromosomally unstable. To test this hypothesis, we perform single cell sequencing on RPE1-p53KO cells 72h after release from 1 or 7 days CDK4/6i-treatment. Figures 1G and H shows that prolonged CDK4/6i treatment leads to increased aneuploidy and heterogeneity 72h after drug washout. This is rescued by combined inhibition of CDK4/6 and mTOR, implying these chromosomal changes are driven by cell enlargement during the G1 arrest. Increased heterogeneity following cell enlargement was also observed in the aneuploid breast cancer line MCF7 (Figure 1I).

In summary, increased cell size causes delays in kinetochore-microtubule attachment and chromosome alignment, however, cells are unable to wait to correct those attachment errors before progressing into anaphase, leading to chromosome segregation errors and aneuploidy.

The mitotic checkpoint, also known as the spindle assembly checkpoint or SAC, is supposed to delay anaphase in the presence of unattached kinetochores^23^ therefore we hypothesised that the SAC may also be weakened in large cells. In support, the duration of mitotic arrest in the presence of either STLC or nocodazole – drugs that prevent kinetochore-microtubule attachment to activate the SAC – was shortened specifically in enlarged CDK4/6i-treated cells (Figures 2A and S2). The SAC produces the mitotic checkpoint complex (MCC), which inhibits the anaphase promote complexes/cyclosome (APC/C) to restrict Cyclin B degradation and prevent mitotic exit^23^. Live cell imaging with an endogenous Cyclin B1-EYFP RPE1 cell line confirmed that enlarged CDK4/6i-treated cells degraded Cyclin B more rapidly during a STLC-induced arrest, consistent with SAC weakening in these cells (Figure 2B).

**Figure 2.**
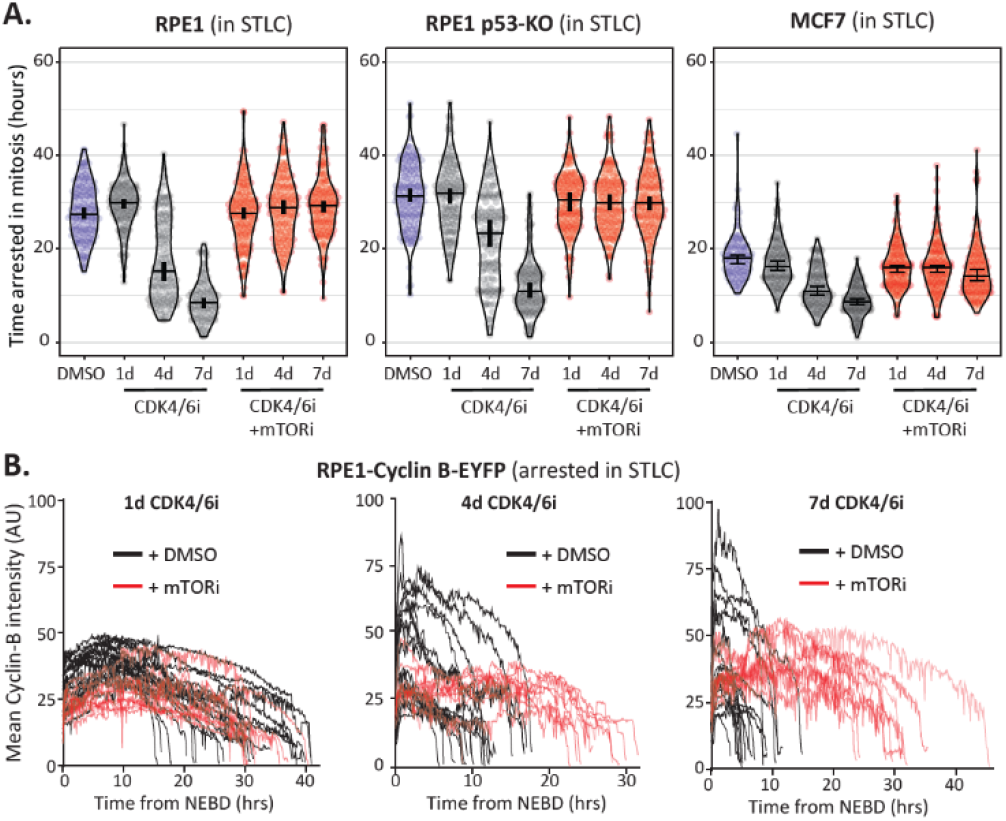
SAC strength is weakened in enlarged CDK4/6i-treated cells. **A-B)** Duration of mitotic arrest (A) and endogenous cyclin B levels (B) in indicated cell lines arrested in mitosis with STLC afte washout from CDK4/6i ± mTORi for indicated durations. In A, measurements were from 150 cells from 3 experiments. Horizontal bars on the violin plots show median, and vertical bars show 95% confidence intervals. In B, mean cyclin B intensities were measured from NEBD until mitotic exit and each individual line represents a single cell. 10 cells per condition from 2 experiments.

In summary, cell enlargement following CDK4/6i-treatment causes delays in kinetochore-microtubule attachment and a weakened mitotic checkpoint, leading to enlarged cells exiting mitosis with chromosome segregation errors. Enlarged CDK4/6i-treated cells also experience earlier replication stress^14,17^, which could potentially affect mitotic fidelity^24^. However, supplementation with excess nucleosides can rescue the replication stress^14^ and increase the percentage of cells entering mitosis (Figure S3A), but this is unable to limit the mitotic delays or SAC weakening (S3B-C). We conclude therefore, that cell size has more direct effects on mitotic fidelity. We next set out to characterise these defects.

### Mechanism of SAC weakening in enlarged cells

The strength of the SAC is determined by relative levels of MCC and APC/C. MCC is produced at unattached kinetochore before being release into the cytoplasm to rapidly inhibit APC/C-CDC20 molecules^23^. We reasoned that the SAC could be weakened in large cells by changes to the relative levels of MCC or APC/C components. Alternatively, if the concentration of MCC and APC/C components remain unchanged, then effective APC/C inhibition may still be compromised because the same number of kinetochores catalyse MCC production, but these inhibitory complexes must then diffuse throughout a larger cytoplasmic volume to inhibit more APC/C molecules.

We therefore assessed the effects of cell size (1-7d CDK4/6i -/+ mTORi) on the levels of MCC and APC components during the G1 arrest or during the subsequent G2/M (Figure 3A). Protein concentrations were normalised in all conditions prior to western blotting so as to provide an estimate of relative changes to cellular concentrations. Figure 3B-C shows that the MCC components MAD2, BUBR1, BUB3 and CDC20 are all enriched in G2/M-arrested MCF7 cells, as expected. However, we observed no clear size-dependent changes to their concentrations under any conditions. Furthermore, the levels of 4 different APC-C subunits were also broadly unaltered by cell cycle stage or cell size (Figure 3D-E). Similar results were observed in RPE1-p53 KO cells (Figure S4), which also displayed considerable size-dependent SAC weakening (Figures 2A and S2). We conclude that cell size does not weaken the SAC by altering the relative concentration of MCC or APC/C subunits. Even though the cytoplasmic levels of MCC components remain unchanged, their recruitment to kinetochores could be perturbed to affect MCC assembly. However, we observed no striking size-dependent changes in the kinetochore levels of MAD1, MAD2, BUBR1 or CDC20 in either nocodazole or STLC-arrested mitotic cells (Figure 3F-G).

**Figure 3.**
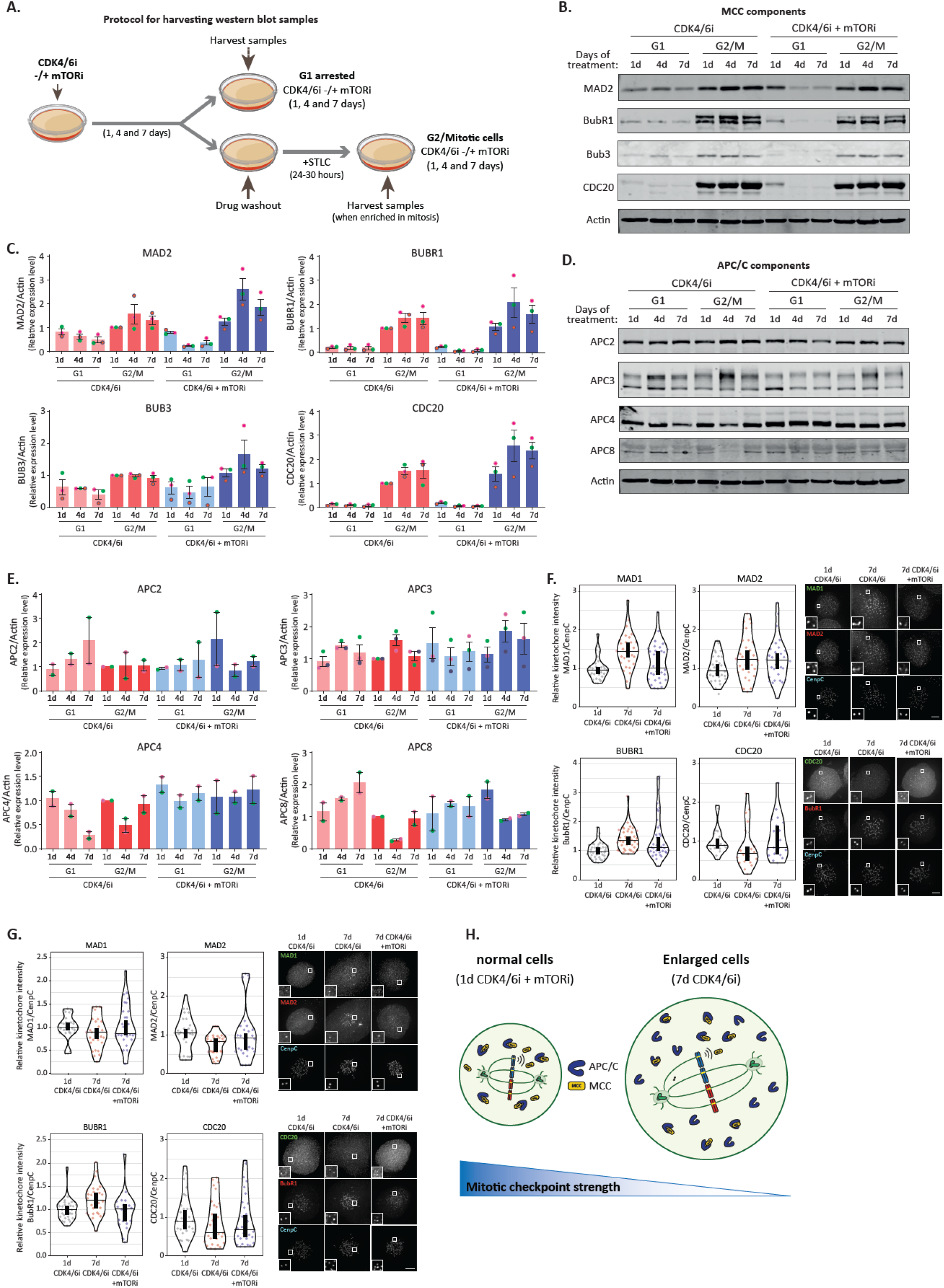
MCC or APC/C concentrations or kinetochore levels are unchanged in enlarged CDK4/6i treated cells. **(A)** Schematic showing protocol used for harvesting samples for immunoblotting. **(B-C)** Immunoblots (B) and quantifications (C) of MCC subunits in G1 arrested and G2/M cells following treatment with CDK4/6i ± mTORi for indicated number of days. Westerns are representative of 3 experiments. **(D-E)** Immunoblots (D) and quantification (E) of APC/C subunits in G1 arrested and G2/M cells following treatment with CDK4/6i ± mTORi for indicated number of days. Westerns are representative of 2-3 experimental repeats. **(F-G)** Quantified kinetochore levels of Mad1, Mad2, BubR1 and Cdc20 in nocodazole-arrested (F) and STLC-arrested MCF7 cells (G). Kinetochore intensities are measured from 30 cells, 3 experimental repeats. Right panels show representative example immunofluorescence images. **(H)** Scheme representing reduced mitotic checkpoint strength in large cells due to increased cytoplasmic volume.

In summary, the level of MCC production at kinetochores is unlikely to be affected by increased cell size. However, the MCC complexes that are produced by kinetochores in large cells must then inhibit more APC/C molecules in a larger cytoplasmic volume, which we predict contributes to SAC weakening (Figure 3H). This is analogous to mechanism proposed for SAC weakening in the early embryonic divisions of large C-elegans embryos^25^

### Mechanism of kinetochore-microtubule attachment defects in enlarged cells

Sister kinetochores are held together by cohesin, which resists the pulling forces exerted by bipolar microtubule attachments. To check for defects in this process in enlarged cells, we measured inter-kinetochore distances on chromosomes that had aligned to the metaphase plate. Figure 4A shows that inter-kinetochore distances were significantly increased in enlarged CDK4/6i-treated cells, suggesting effective microtubule attachment but impaired sister chromatid cohesion. Cohesion is gradually lost from aligned kinetochores under tension due to a process known as cohesion fatigue^26^. However, the reduced cohesion was not due to the mitotic delays in enlarged cells because inter-kinetochore distances were also increased 30 min after release from R03306 to synchronise mitotic entry (Figure S5).

**Figure 4.**
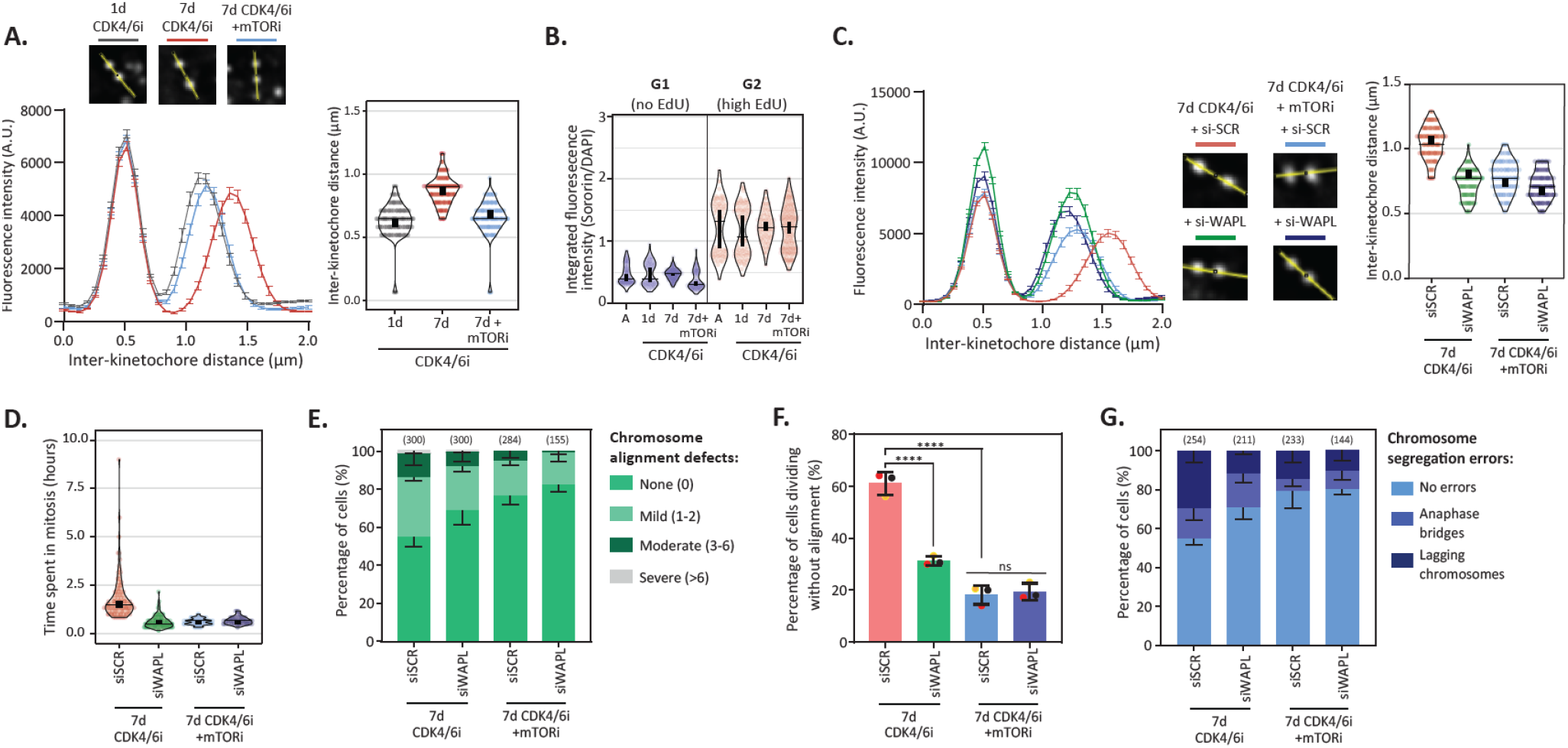
Cohesion defects in large CDK4/6i treated cells contribute to chromosome segregation errors. **A)** Inter-kinetochore distances in MCF7 cells following treatment with CDK4/6 ± mTORi for indicated times. Left panel shows CenpC line plots showing the inter-kinetochore distances. The two peaks indicate the kinetochore pairs. 5 kinetochore pairs were analysed per cell, for a total of 10 cells per experiment. Graphs represent the fluorescence intensities (±SEM) from 3 independent experiments. Top panel shows representative kinetochore pairs for each condition. Right panel shows the measured inter-kinetochore distances (μm). **(B)** Levels of sororin in MCF7 cells during G1 and G2 phase following drug washout from CDK4/6 ± mTORi. **(C)** Inter-kinetochore distances in 7d CDK4/6i ± mTORi treated cells after siRNA depletion of WAPL. Left panel shows CenpC line plots showing the inter-kinetochore distances. Middle panel shows representative kinetochore pairs for each condition and right panel shows the measured inter-kinetochore distances (μm). **(D-G)** Effects of WAPL depletion on mitotic errors in large 7d CDK4/6i ± mTORi treated cells. Violin plots in (D) shows mitotic duration. Measurements are from 100 cells from 2 experimental repeats. Graphs in panel (E) and (G) show mean frequencies of chromosome misalignments and chromosome segregation errors, respectively (±SEM; 3 experiments, with total number of cells quantified per condition indicated above each bar). Panel (F) shows percentage of cells dividing before chromosome alignment is complete. Statistical significance was determined by Chi-square test (****p<0.0001). Data shows mean ± SEM from three repeats. For violin plots in A-D, horizontal bars show median, and vertical bars show 95% confidence intervals. which can be used for statistical comparison of different conditions (see Materials and Methods).

Sister chromatid cohesion is established during DNA replication and maintained thereafter by sororin which antagonises the action of the cohesin-release factor WAPL^27^. Chromatin-bound sororin, which can be used as a marker for cohesive cohesion^28^, was specifically elevated following DNA replication, as expected, but this was unaltered in enlarged CDK4/6i-treated cells (Figure 4B). Considering we observed no obvious defect in cohesin establishment in enlarged cells, we hypothesised that cohesin may instead be prematurely removed by WAPL during mitosis. In agreement, WAPL knockdown specifically rescued inter-kinetochore distances in enlarged CDK4/6i-treated cells (Figure 4C). This was also sufficient to completely rescue the mitotic delays and partially rescue the chromosome alignment defects and segregation errors (Figures 4D-G). Therefore, WAPL-mediated release of cohesin causes mitotic delays, kinetochore-microtubule attachment errors and chromosome missegregations in enlarged CDK4/6i-treated cells.

In human cells, WAPL is known to remove cohesin from chromosomes during early mitosis^29,30^ when its antagonist sororin is displaced from cohesin by mitotic kinases^31,32^. Importantly, centromeric cohesion is preserved by shugoshin (Sgo1) which competes with WAPL for cohesin-binding and recruits PP2A to prevent sororin phosphorylation^27,33^. Together, this maintains cohesion at centromeres, where it is needed to resist pulling forces exerted by kinetochore-microtubules. Immunofluorescence imaging of nocodazole-arrested cells showed that Sgo1 localisation was defective in enlarged CDK4/6i-treated cells. Total levels of Sgo1 at the centromere/kinetochore were largely unaffected by cell size, but Sgo1 was relocalised to kinetochore proximal region (Figure 5A,B). Sgo1 binds directly to cohesin, therefore Sgo1 relocalisation could be a cause and/or consequence of cohesin loss from centromeres. To investigate this further, we recovered Sgo1 at centromeres by fusing to CenpB (CB-Sgo1), which was sufficient to also rescue cohesion in enlarged cells, as judged by a reduction in inter-kinetochore distances (Figure 5C and S6). Similarly, rescuing cohesion by WAPL depletion also recovered Sgo1 localisation to centromeres (Figure 5D). Therefore, the ability of Sgo1 to antagonise WAPL and protect cohesion at centromeres is impaired in enlarged CDK4/6i-treated cells. This causes premature loss of cohesion and chromosome segregation errors.

**Figure 5.**
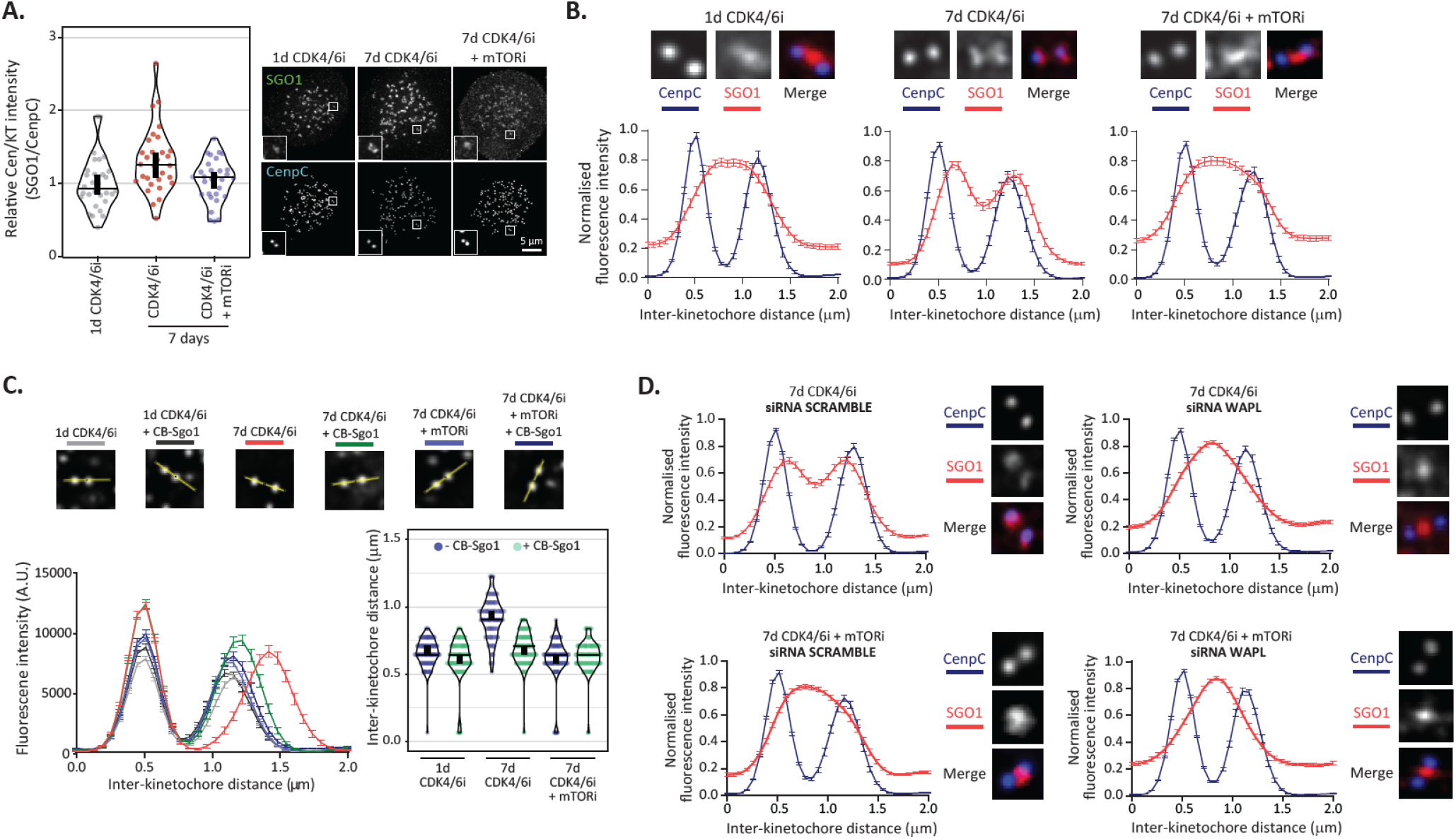
Sgo1 relocalisation contributes to cohesion defects in enlarged CDK4/6i treated cells. **(A)** Levels of Sgo1 at unattached kinetochores in nocodazole arrested MCF7 cells following treatment with CDK4/6i ± mTORi, as indicated. Kinetochore intensities are measured from 30 cells, 3 experimental repeats. Right panel shows representative example immunofluorescence images. Insets show magnifications of the outlined regions. Scale bars: 5µm, Inset size: 1.5µm. **(B)** Line plots of Sgo1 and CenpC localisation across the kinetochore:centromere axis in cells treated as in A. 5 kinetochore pairs were analysed per cell, for a total of 10 cells per condition. Graphs represent the mean intensities (±SEM) from 3 experiments. Intensity is normalized to the maximum signal present in each channel in each experiment. Top panel shows representative localisation images for Sgo1 and CenpC for each condition. **(C)** Interkinetochore distances on metaphase aligned kinetochores in CDK4/6 ± mTORi treated cells that were transfected with/without CB-Sgo1. Right panel shows the measured inter-kinetochore distances (μm) and top panel shows representative images for each condition.5 kinetochore pairs were analysed per cell, for a total of 10 cells per condition. Graphs represent mean of fluorescence intensities from 2 experiments (±SEM). MG132 was added to these cells to prevent mitotic exit and arrest cells in metaphase. **(D)** Line plots of Sgo1 and CenpC localisation in 7d CDK4/6 ± mTORi treated cells following WAPL depletion. Right panels next to each individual line plot indicate representative localisation images for each condition. 5 kinetochore pairs were analysed per cell, for a total of 10 cells per condition from 2 experiments. Graphs show mean intensities (±SEM). Intensity is normalised to the maximum signal in each channel in each experiment.

## Discussion

This work reveals how increases to cell size following CDK4/6i treatment can have deleterious effects on the segregation of chromosomes during mitosis. Enlarged cells have a weakened mitotic checkpoint response, most likely because the same number of kinetochores must catalyse more MCC to inhibit the additional APC/C molecules present within a larger cytoplasmic volume. When all kinetochores are signalling, cells can delay anaphase for a few hours, but when just a few unattached kinetochores remain, these are most likely insufficient to halt mitotic progression. This is particularly problematic for large cells because Sgo1 is also delocalised from centromeres causing weakened sister chromatid cohesion and kinetochore-microtubule attachment delays. The inability to wait to properly attach those kinetochores leads to chromosome segregation errors and aneuploidy in enlarged CDK4/6i-treated cells.

This has important implications for the emergence of CDK4/6i resistance in advanced HR+/HER2-breast cancer patients. In this setting, p53 mutations are some of the most common genetic alterations associated with acquired and intrinsic resistance, and this was recently linked to the inability of p53 deficient cells to enter senescence following CDK4/6i-treatment^19–22^. We hypothesise that p53-deficient cells that evade senescence can use their enlarged size to induce chromosome segregation errors before selecting for karyotypes and/or genotypes that offer proliferative advantages. In this model, oncogenic mutations, which are also frequently associated with resistance^19–22^, are predicted to drive chromosome missegregations by increasing cell size during the CDK4/6i arrest. Furthermore, preventing increases to cell size by inhibiting signalling to mTOR should limit resistance by rescuing chromosome segregation errors, as we observe following mTORi in MCF7 and RPE1 cells. This could help to explain why the PI3K inhibitor inavolisib delays the onset of resistance in PIK3CA-mutated breast cancer patients treated with palbociclib and fulvestrant^34^. It will be important in future to address whether increases to cell size are observed in patient tumours, and if so, whether these enlarged cells renter the cell cycle to cause chromosome segregation errors during the frequent CDK4/6i holiday periods. If they do, then the molecular basis for these mitotic errors could perhaps be targeted to limit drug-resistance.

The work we have presented on CDK4/6i could be widely relevant for other anti-cancer treatments that also increase cell size. For example, a growing body of evidence indicates that chemotherapeutics that damage DNA to indirectly arrest the cell cycle also induce cell overgrowth during this arrest^35–37^. Therefore, enlarged cells that evade therapy-induced senescence in these contexts, or even cells that enter senescence but then later escape, could be similarly primed to experience chromosome segregation and aneuploidy because of their increased size. This may help to explain why cells that escape from therapy-induced senescence can drive such aggressive tumourigenesis^38–40^.

Finally, our work could also be relevant in physiological contexts in the body when cells become enlarged. One good example is during aging of hematopoietic stem cells (HSCs) where increased cell size contributes to decreased replicative potential and functional decline^41^. Aneuploidy also impairs HSC fitness^42^, therefore increased cell size could contribute to functional decline of aged HSC by inducing aneuploidy. If increased size causes permanent genetic changes during the aging process, this might also explain why rapamycin is able to prevent the functional decline of aged HSCs and increase longevity in mice, but still be unable to restore HSC function once cells have already become enlarged^43,44^. It will be important in future to determine whether increased cell size can drive age-associated aneuploidies in HSCs and in other cell types and tissues.

## Acknowledgements

We thank staff at the Dundee Imaging Facility. This work was funded by a Ninewells Cancer Campaign Doctoral Training Programme which funded A.P., and a Cancer Research UK Programme Foundation Award to A.T.S (C47320/A21229).

## Author Contributions

A.T.S conceived and supervised the project. A.P. performed and analysed all experiments, except for the single cell sequencing which was performed by M.L. under the supervision of F.F. A.T.S. wrote the original manuscript draught with review and editing by A.P., who also generated all figures.

## Materials and Methods

### Cell culture and reagents

hTERT-RPE1 cells and the human ER+/HER2-breast cancer cell line MCF7 were purchased from ATCC. The RPE1 Cyclin B1-eYFP cells have been described previously^45^. All cells were cultured at 37°C with 5% CO2 in DMEM (Thermo Fisher Scientific, Gibco, 41966029) supplemented with 9% FBS (Sigma Aldrich, F7524) and 50 μg/ml penicillin/streptomycin (Thermo Fisher Scientific, Gibco, 15070063). Every 4–8 weeks, cells were screened to ensure a mycoplasma free culture.

The following drugs were used in this study (including final concentrations used throughout this study): palbociclib (PD-0332991, hydrochloride salt, LC Laboratories, P-7788; 1μM for MCF7/MCF7-Hist1H4C-GFP and 1.25μM for RPE1/RPE1 p53-KO), PF-05212384 (gedatolisib Sigma, PZ0281; 7.5nM for MCF7/MCF7-Hist1H4C-GFP and 30nM for RPE1/RPE1 p53-KO), nutlin-3a (Sigma, SML0580; 5µM), S-Trityl-L-cysteine (STLC; Sigma Aldrich, #164739; 10μM), nocodazole (Sigma Aldrich, 487928; 3.3μM), RO-3306 (Tocris, 4181; 10μM), MG132 (Sigma Aldrich, 474787; 10μM), EmbryoMax Nucleosides 100X (Merck, ES-008-D; 1:100 dilution). The vsv-CENP-B-Sgo1-mCherry plasmid was described previously^46^.

### Generation of cell lines

For generating the MCF7 Hist1H4C-GFP cell line, the endogenous Hist1H4C in MCF7 cells was tagged with TagGFP2 by using the CRISPaint method^47^. Briefly, cells were transiently transfected with pCas9-mCherry-Frame1 selector, pCas9-mCherry-Hist1H4C target selector and pCRISPaint-TagGFP2-PuroR donor plasmid at a ratio of respectively 1:1:2, using Fugene HD (Promega) according to the manufacturer’s instructions. After two days from transfection, cells were selected using puromycin for four weeks. The p53 knockout cell line in RPE1 was generated by CRISPR/Cas9 using a gRNA targeting exon 4 of p53 (ACCAGCAGCTCCTACACCGG) followed by selection in 5µM nutlin-3A, as described previously (Crozier et al., 2022; Crozier et al., 2023).

### Gene knockdown

The siRNAs used in this study were siWAPL (5’-GAGAGAUGUUUACGAGUUU-3’, UU overhang) and siScramble (control siRNA: 5'-AAGCGCGCTTTGTAGGATTCG-3’, UU overhang).

All synthesised siRNAs (Sigma-Aldrich) were used at 20 nM final concentration. All siRNAs were transfected using Lipofectamine® RNAiMAX Transfection Reagent (Thermo Fisher) according to the manufacturer’s instructions. After 24 h of knockdown, cells were treated with palbociclib -/+ PF-05212384 for indicated days. Cells were then released from the drug into full growth media, and when appropriate, into STLC for live-cell assays or MG132 for fixed analysis.

Cell volume analysis: For cell volume analysis, cells were plated into 6-well plates at a density of 30,000 cells per well then treated with drugs immediately, and incubated at 37 °C with 5% CO_2_. At 24 h intervals, cells were trypsinised, stained with acridine orange and DAPI, and cell diameters were measured using a Chemometec NC-3000 Nucleocounter. Histograms of cell diameter were imported into Flowing Software version 5.2.1 and mean cell diameter of each condition was taken to calculate volume using 4/3 πr^3^.

### Immunofluorescence

Cells were plated at low density on High Precision 1.5H 12-mm coverslips (Marienfeld) and fixed for 10 min with 4% paraformaldehyde dissolved in PBS or pre-extracted with 0.1% Triton X-100 in PEM (100 mM PIPES, pH 6.8, 1 mM MgCl2 and 5 mM EGTA) for 1 min before addition of 4% PFA for 10 min. After fixation, coverslips were washed three times in PBS and then blocked with 3% BSA dissolved in PBS with 0.5% Triton X-100 for 30 min. Coverslips were then incubated with primary antibodies overnight at 4°C, prior to washing with PBS and incubation with secondary antibodies and DAPI (4,6-diamidino2-phenylindole, Thermo Fisher; 1 μg/ml) for 2–4 h at room temperature in the dark. Coverslips were washed with PBS and mounted onto slides with ProLong Gold Antifade (Thermo Fisher Scientific, P36930). Images were acquired on either a Zeiss Axio Observer using a Plan-apochromat 20×/0.8 NA Air objective, a Deltavision Core or Elite system with a 100×/1.40 NA U Plan S Apochromat objectives, or a Nikon Ti2-E Eclipse with a Kinetix camera (Teledyne Photometrics) at 4×4 binning using a CFI Plan Apochromat λD 20x/0.8 NA air objective. For cells transfected with CB-Sgo1 plasmid, mitotic cells arrested in metaphase were selected for imaging based on good expression of mCherry on the kinetochore/centromere. All immunofluorescence images displayed are maximum intensity projections of deconvolved stacks and were chosen to closely represent the median quantified in the data. Figure panels were created using Omero (http://openmicroscopy.org).

The primary antibodies were used at the final dilutions as indicated: guinea pig anti-CENP-C (PD030 from Caltag + Medsystems, 1:5000), mouse anti-BUB1 (ab54893 from Abcam, 1:400), rabbit anti-BUBR1 (A300-386A from Bethyl, 1:1,000), mouse anti-MAD1 (MABE867 from Millipore, 1:1000), mouse anti-CDC20 (sc13162 from Santa Cruz, 1:1000) rabbit anti-MAD2 (custom polyclonal, 1:2000), rabbit anti-SKAP (HPA042027 from Atlas Antibodies, 1:1000), rabbit anti-sororin (ab192237 from Abcam, 1:1000) and rabbit mCherry (GTX128508 from Genetex, 1:1000). For EdU staining, a base click EdU staining kit was used (Sigma, BCK-EDU488), as per manufacturer’s instructions. The secondary antibodies used were highly cross-absorbed goat anti-mouse Alexa Fluor 488 (A-11029), goat anti-rabbit Alexa Fluor 568 (A-11036) and goat anti-guinea pig Alexa Fluor 647 (A-21450) all used at 1:1000 (Thermo Fisher).

### Time-lapse imaging

For brightfield movies to measure mitotic duration or SAC strength, cells were plated in a 24-well plate and treated with palbociclib -/+ PF-05212384 for different lengths of time (1, 4 and 7 days). Following drug washout, cells were incubated with full growth media or in the presence of nocodazole or STLC in a heated chamber (37°C and 5% CO2) and imaged with brightfield microscopy with a 10×/0.5 NA air objective using a Zeiss Axio Observer 7 with a CMOS ORCA flash 4.0 camera at 4 × 4 binning with images taken every 10 minutes apart. For each condition, time of entry into mitosis (defined by rounding up of the cell) and time of mitotic exit (defined by cell division or cells flattening down in presence of nocodazole/STLC) were recorded for 50 random cells selected at the beginning of the time lapse.

### Chromosome alignment and segregation assay

To observe live chromosome alignment and determine mitotic cell fates, MCF7-HIST1H4C-tagGFP cells were plated and treated in 8-well chamber slides (ibidi). Following drug washout, cells were imaged every 5 min for 72 h with a 40×/1.3 oil immersion objective using a Zeiss Axio Observer 7 with a CMOS Orca flash 4.0 camera at 4 × 4 binning and 10 z-stacks with a step size of 1.50 μm. Selected cells were scored based on the whether mitotic cells underwent division or cell death following chromosome alignment or not.

To observe chromosome alignment in fixed-cell experiments, cells were released from palbociclib -/+ PF-05212384 after treatment and released in full-growth media. Cells were then incubated for 16-24 h before addition of MG132 for 30 min, to arrest cells in metaphase and subsequently fixed and stained as described above and imaged on a Zeiss Axio Observer with a CMOS Orca flash 4.0 camera at 4 × 4 binning, using a Plan-apochromat 20×/0.4 air objective. Cells were scored based on the number of misaligned chromosomes as following: aligned (0 misaligned chromosomes, with a visible metaphase plate), mild (1–2), moderate (3–6) and severe (> 6 misaligned chromosomes). To observe chromosome segregation defects in fixed-cell experiments, following drug washout from palbociclib -/+ PF-05212384, cells were released in full growth media and fixed 24-30 h following washout to allow cells to enter anaphase. Anaphase cells were scored based on the type of chromosome segregation defects (no errors, anaphase bridges or lagging chromosomes).

To measure the number of unattached kinetochores, cells were released in full-growth media following washout from palbociclib -/+ PF-05212384. Prior to mitotic entry, cells were treated with RO-3306 for 2 h to synchronise cells at the G2/M boundary. Cells were then washed three times and incubated for further 15 min with full-growth media before addition of MG132 to prevent mitotic exit. After 30 minutes from MG132 addition, cells were fixed and stained as described above. The number of unattached kinetochores on the mitotic spindle was quantified by counting the number of kinetochores positive for MAD1 recruitment.

### Western blotting

As shown in Fig 3A, cells were treated with palbociclib -/+ PF-05212384 for 1, 4 and 7 days following which cells were either harvested during the drug-induced arrest or released into full-growth media supplemented with STLC following drug washout to prevent cells from exiting mitosis and harvested 24-30 h after when cells were enriched in mitosis. Protein lysates were prepared by scraping cells into 2X SDS buffer (100mM Tris, pH 6.8, 4% SDS, 20% glycerol, 0.2% bromophenol blue and protease inhibitor cocktail), sonicated with a Cole-Parmer ultrasonic processor (20% Amp, 15 sec pulse), followed by DC protein assay (Biorad) to estimate the concentration of each sample, after which 2-mercaptoethanol was added at a final concentration of 10%. Samples were boiled, and equal concentrations of protein were loaded and separated on 12% or 10% SDS-PAGE gels and transferred onto 0.45µm nitrocellulose membranes (Amersham Protran Premium). After transfer, blots were blocked in 5% milk in TBS with 0.1% Tween 20 (TBS-T) and incubated overnight at 4°C in primary antibodies. Membranes were then washed three times in TBS-T, incubated in IRDye secondary antibody for 1h, and washed 3 further times prior to visualisation on a LI-COR Odyssey CLx system.

The following primary antibodies (all diluted in 5% non-fat milk in TBST) were used at the final dilutions indicated: rabbit anti-MAD2 (custom polyclonal, 1:1000), rabbit anti-BUBR1 (Bethyl, A300-386A, 1:2000), mouse anti-BUB3 (BD Transduction, 611730, 1:1000), mouse anti-CDC20 (1:1000), mouse anti-APC3 (BD Transduction, 610454, 1:1000), rabbit anti-APC4 (Bethyl, A301-176A, 1:1000), rabbit anti-APC2 (Cell Signalling, 12301S, 1:1000), rabbit anti-APC8 (Bethyl, A301-181A, 1:1000), rabbit anti-Actin (Sigma Aldrich, A2066, 1:2500). The secondary antibodies used were IRDye 800CW goat anti-mouse IgG (LICORbio, 926-32210, 1:15,000) and IRDye 800CW donkey anti-rabbit IgG (LICORbio, 926-32213, 1:15,000).

### Single cell library preparation and sequencing

Cells previously frozen in DMSO were thawed, pelleted by centrifugation, and resuspended in 1 mL of lysis buffer per 1 × 10^6^ cells. The buffer (10 mM Tris-HCl pH 7.4, 154 mM NaCl, 0.2% BSA, 0.1% NP-40, 1 mM CaCl_2_, 0.5 mM MgCl_2_ in ultra-pure water) was used to selectively lyse the cell membrane and release intact nuclei. Nuclei were stained with propidium iodide (10 µg/mL) and Hoechst 33258 (10 µg/mL) to allow sorting based on cell cycle state and viability. Single nuclei were dry-sorted into 384-well plates using a MoFlo Astrios sorter (Beckman Coulter) and frozen at –80°C. Libraries were prepared using the cellenONE® F1.1/F1.4 v2.0 nanodispenser (Cellenion), which performs DNA deproteinization and shearing, followed by ligation of dual 10 bp barcoded adapters in each well. The libraries were then pooled, amplified, and purified using magnetic beads, as described previously^48^. Sequencing was performed on an Illumina NextSeq 2000 platform generating 77 bp single-end reads. Sequencing data were demultiplexed into FASTQ files using bcl2fastq (Illumina).

### AneuFinder analysis and scoring

Sequencing reads were aligned to the GRCh38/hg38 human genome assembly using Bowtie2 (v2.2.4)^49^, and duplicate reads were marked using BamUtil (v1.0.3)^50^. Copy number analysis was performed with AneuFinder (v4.3.3) following the method in Bakker et al.^51^. The analysis included GC correction and blacklisting of artifact regions identified from euploid controls. Copy number calling was conducted using the dnacopy and edivisive algorithms on variable-length bins averaging 1 Mb with 500 kb steps. Libraries with less than 10 reads per chromosome copy per bin were excluded. Aneuploidy scores were calculated as weighted averages of absolute copy number deviations from euploid state. Heterogeneity scores were determined using the original approach from Bakker et al.^51^, based on copy number variation. For samples containing tetraploid libraries, heterogeneity scores were calculated by first determining the copy number differences relative to the median copy number profile, which reflects the normal copy number state of the cell line (3-somy for chromosome 10q, 4-somy for chromosome 12p, and 2-somy for all other chromosomes). For tetraploid libraries, differences were calculated relative to the doubled median copy number profile.

### Image quantification and statistical analysis

For quantification of kinetochore protein levels, 10 mitotic cells (nocodazole arrested) were randomly selected, and images were acquired on DelvaVision Elite or Core system as mentioned above. Images of similarly stained experiments were acquired with identical illumination settings and analysed on Fiji, Image J software using an ImageJ macro, as described previously^52^. Briefly, the macro performs a threshold and selection of all the kinetochores, using the DAPI and CenpC channels. To generate kinetochore masks, the macro applied a convolution filter to the CenpC channel and perform a threshold selection. The resulting masks were then increased by 1 pixel (to ensure complete kinetochore selection). These masks were used to calculate the relative mean kinetochore intensities of indicated proteins. Fluorescence intensities at kinetochores were normalised to the kinetochore marker, CenpC.

For quantification of Sgo1 localization, The CenpC channel was used to choose 5 random kinetochore pairs per cell that lie on the same 0.2 μm Z-plane. A line was then drawn through the kinetochore pairs (using ImageJ), with the first CenpC kinetochore peak at 0.5 µm from the start of the line. An ImageJ macro (created by Kees Straatman, University of Leicester and modified by Balaji Ramalingam, University of Dundee) was used to simultaneously measure the intensities in each channel across the line. The signal from the five kinetochore pairs was averaged and normalized to the maximum signal in each channel. Inter-kinetochore distance for each kinetochore pair was measured as the distance between the two peaks from the intensity profiles from the CenpC channel.

The single-cell FUCCI profiles were generated manually by analysing RPE1-FUCCI movies. A total of 50 red cells were randomly selected and marked at the beginning of the movie. The time points in which the FUCCI cells change colour was recorded to determine the time spent in each phase of the first cell cycle following release from CDK4/6 inhibition. RPE-FUCCI cells were always imaged with the same illumination settings and all images were placed on the same scale prior to analysis to ensure that the red/yellow/green cut-offs were reproducibly calculated between experiments. Mitotic entry was timed based on the visualization of typical mitotic cell rounding and loss of nuclear mAG-geminin signal.

Sororin intensities were calculated by first using the DAPI channel to generate an ROI overlay in ImageJ. This overlay was then applied to the DAPI, EdU and Sororin channels and used to measure the mean grey value of each individual ROI. An area outside any ROI was then designated as background and its mean grey value in either channel was also calculated. This background value was then subtracted from all ROI values from the corresponding channel. Since cells were release from palbociclib into EdU and STLC, those cells lacking EdU were designated as G1 cells and cells expressing high EdU were designated as G2 cells. Their corresponding sororin values were used to calculate integrated fluorescence intensity.

Violin plots were produced using PlotsOfData — https://huygens.science.uva.nl/PlotsOfData/^53^. This allows the spread of data to be accurately visualised along with the 95% confidence intervals (thick vertical bars) calculated around the median (thin horizontal lines). This representation allows the statistical comparison between all treatments and time points because when the vertical bar of one condition does not overlap with one in another condition, the difference between the medians is statistically significant (P < 0.05). Chi-square test was performed to compare the different experimental groups in Fig 1E and Fig 4F (using GraphPad Prism 10 software). All the other plots were generated using GraphPad Prism 10.

## Declaration of interests

The authors declare no competing interests

## Notes

### Competing Interest Statement

The authors have declared no competing interest.

### Summary of Updates

The text in the results section was modified slightly

