## Supplementary Figures for "Cell enlargement causes mitotic errors and aneuploidy in cells that evade senescence after CDK4/6 inhibition"

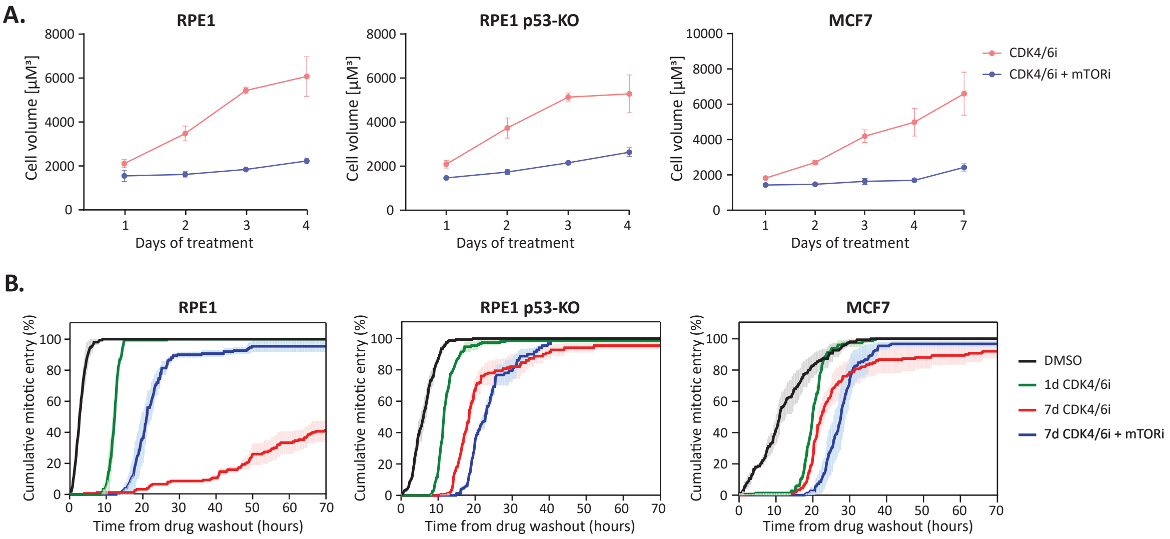


**Figure S1. Cell growth and cumulative mitotic entry following treatment with CDK4/6 inhibitors. A)** Cell volume assays following treatment with palbociclib (CDK4/6i) for indicated time ± PF-05212384 (mTORi: 30nM for RPE1 and 7.5nM for MCF7). Graphs show mean data ± SD from 3 repeats. **B)** Time-lapse analysis of cumulative mitotic entry after washout from indicated drug treatments. Graph show mean ± SEM from 150 cells from 3 experiments.

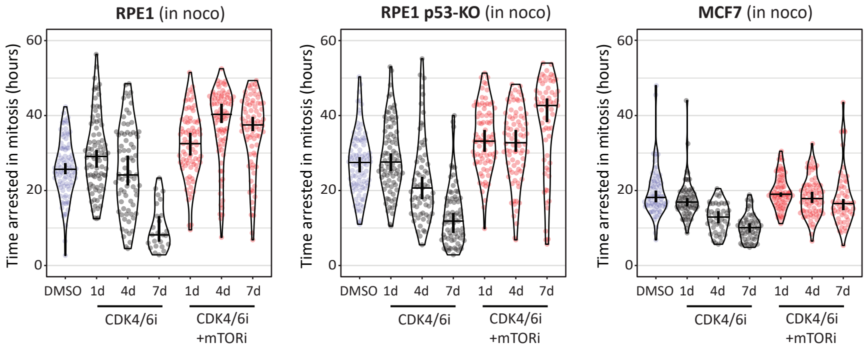


**Figure S2. Mitotic arrest duration in nocodazole is reduced in enlarged CDK4/6i-treated cells.** Duration of mitotic arrest in nocodazole-arrested cells following drug washout from CDK4/6i ± mTORi. Measurements were performed in 150 cells from 3 experiments. Horizontal bars on the violin plots show median, and vertical bars show 95% confidence intervals.

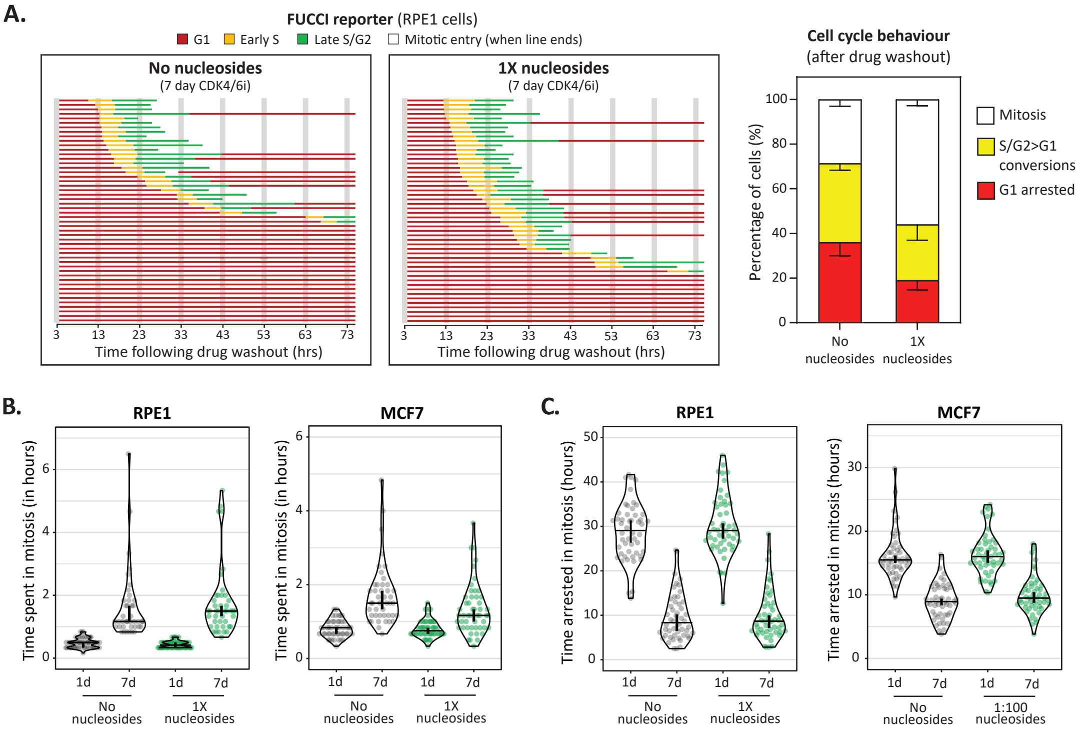


**Figure S3. The mitotic defects in enlarged CDK4/6i-treated cells are not rescued by exogenous nucleoside addition. A)** Cell cycle profile of individual RPE1-FUCCI cells (each bar represents one cell) after washout from 7 days CDK4/6i treatment. Exogenous nucleosides were added 24 h before drug washout and released back into full growth media containing nucleosides and STLC. Right panel shows quantification of cell cycle fates from the single-cell profiles displayed. A total of 150 cells were analysed at random from 3 experiments. **B-C)** Effects of exogenous nucleosides on (B) mitotic delays and (C) length of mitotic arrest in STLC following drug washout from CDK4/6i ± mTORi for indicated times. Measurements were performed in 100 cells from 2 experiments. Horizontal bars on the violin plots show median, and vertical bars show 95% confidence intervals.

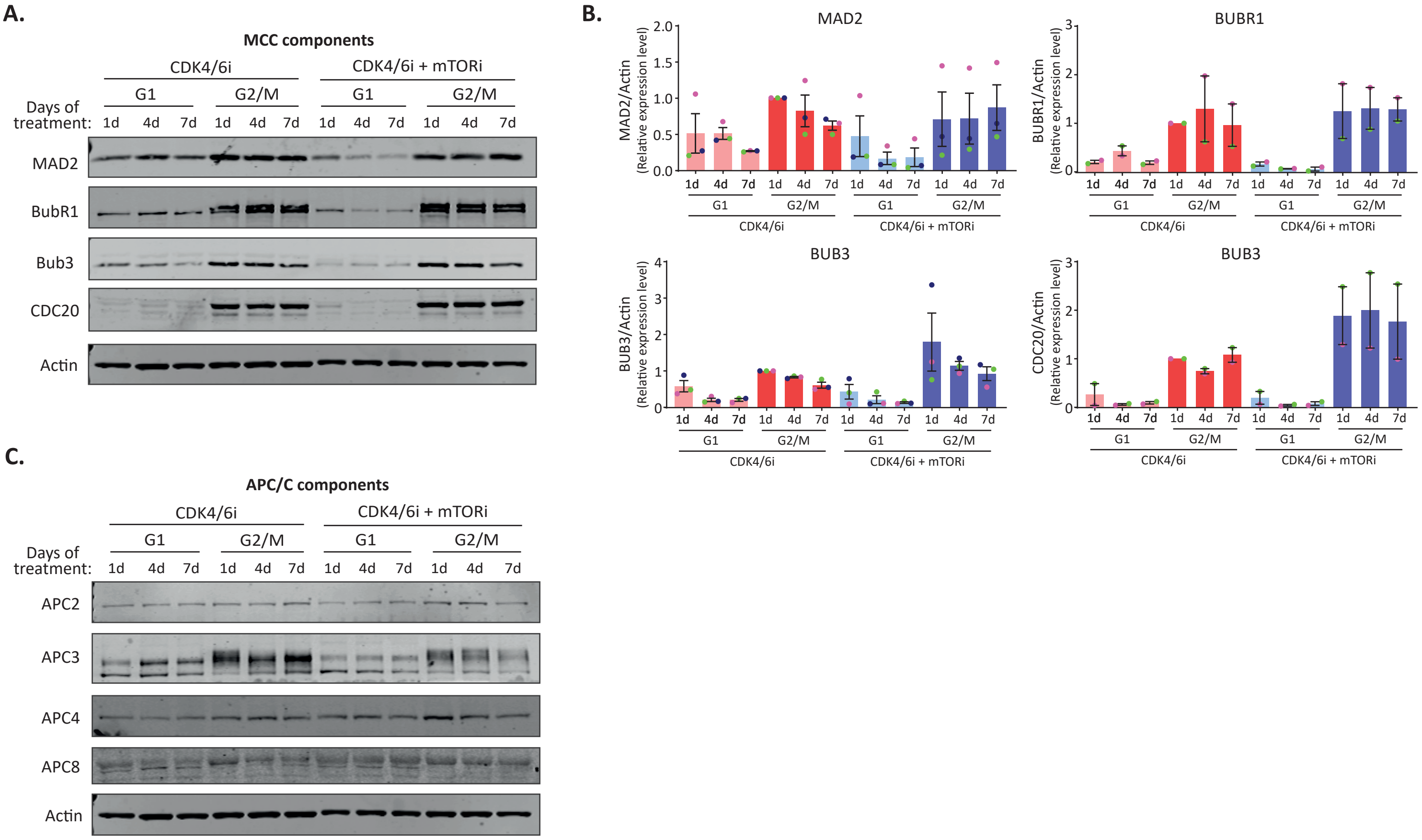


**Figure S4.Levels of MCC and APC/C subunits are unchanged in enlarged CDK4/6i-treated RPE1 p53-KO cells. A-B)** Immunoblot (A) and quantifications (B) of MCC subunits in G1 arrested and G2/M cells following treatment with CDK4/6i ± mTORi for indicated number of days. Westerns are representative of 2-3 experimental repeats. **C)** Immunoblot of APC/C subunits treated as in A.

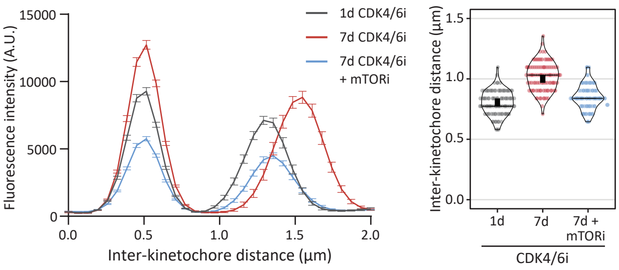


**Figure S5. Cohesion defects in enlarged CDK4/6i treated cells do not result from mitotic delays.** Inter-kinetochore distances in MCF7 cells following treatment with CDK4/6 ± mTORi. Cells were synchronised in G2/M with 10µM RO3306 and released in MG132 for 30min. Left panel shows line plots showing the inter-kinetochore distances. The two peaks indicate the kinetochore pairs. 5 kinetochore pairs were analysed per cell, for a total of 10 cells per experiment. Graphs represent the fluorescence intensities (±SEM) from 2 independent experiments. Right panel shows the measured inter-kinetochore distances (µm).

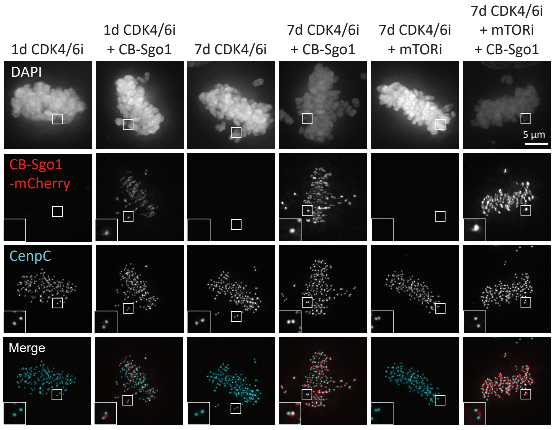


**Figure S6. CenpB-Sgo1 expression at metaphase centromeres. I**mmunofluorescence analysis showing mCherry-CenpB-Sgo1 expression at metaphase centromeres.
